# Symmetry-breaking flow transition as an under-recognized hemodynamic factor in valve-associated thrombosis

**DOI:** 10.64898/2026.07.20.739554

**Authors:** Yuxin Chen, Daniele Vigolo

**Author notes:** Electronic mail.

## Abstract

Venous thrombosis commonly develops near venous valve pockets, where disturbed hemodynamics and reduced washout create spatially heterogeneous flow environments. Conventional descriptions of this region emphasize continuously varying quantities such as velocity, shear stress, recirculation, and residence time, but do not address whether post-valve flow can transition between distinct organized states. Here, we combine computational and experimental vein-on-a-chip approaches to resolve Reynolds-number-dependent post-valve flow organization and its downstream transport consequences. In a geometrically symmetric rigid valve, increasing Reynolds number triggered a symmetry-breaking transition from a centered jet to a persistent asymmetric flow state, with repeatable separation between the increasing- and decreasing-Reynolds-number branches under transient forcing. Valve compliance delayed this transition by increasing the effective opening and reducing local inertial forcing, whereas a small bilateral geometric bias promoted earlier branch selection. Fluid–structure interaction further showed that, after symmetry breaking, unequal hydrodynamic loading generated unequal leaflet deformation, providing a mechanism for structural reinforcement of the selected flow state. The Reynolds-number-dependent ordering was reproduced experimentally and remained detectable in whole blood. Downstream transport of RBC-scale particles exhibited a delayed but persistent lateral redistribution relative to the velocity-field response. These results identify post-valve symmetry breaking as a flow-state transition regulated by local inertial forcing, valve mechanics, and geometric bias. Venous-valve hemodynamics can therefore transition between distinct flow states, providing a physical basis for how small mechanical or geometric differences can generate persistent transport heterogeneity relevant to thrombosis.

## I. INTRODUCTION

Deep vein thrombosis (DVT) frequently develops in the vicinity of venous valve pockets, where reduced flow and limited exchange with the main lumen create conditions favorable to local blood retention^1–3^. Prolonged residence within the valve sinus can promote hypoxia, endothelial activation, leukocyte recruitment, platelet participation, and activation of coagulation pathways^2–7^. These processes place the local hemodynamic environment at the center of thrombus initiation alongside the classical contributions of stasis, vessel-wall dysfunction, and changes in blood composition^3,8–10^.

Flow within the valve region is commonly described through quantities such as velocity, shear rate, recirculation, and residence time. These measures resolve local hemodynamic disturbances but do not directly describe whether the post-valve flow field can reorganise between qualitatively distinct states. Such flow-state changes are well established in confined internal flows and provide a useful framework for considering the global organization of the downstream valve flow.

In symmetric sudden expansions, increasing Reynolds number destabilises a centered laminar jet and promotes mirror-related asymmetric states^11,12^. The onset and structure of these states depend on expansion ratio and confinement^13^, and experiments have identified steady symmetry breaking followed by time-dependent behavior at higher flow rates^14^. These studies show that geometrically symmetric systems can support persistent asymmetric flow and that small perturbations can influence lateral state selection.

Venous valves share several hydrodynamic features with confined expansion flows, including a restricted opening, downstream expansion, separated recirculation regions, and a high-velocity jet. Their deformability introduces an additional mechanical degree of freedom because the effective opening changes under fluid loading. Human lower-extremity veins contain bicuspid valves with substantial anatomical variation in valve position and local venous geometry^15^. Microfluidic valve models have further shown that leaflet stiffness and geometry influence valve opening, local hemodynamics, particle transport, and thrombotic localization^16,17^, while endothelialized vein-on-a-chip systems have demonstrated mechanically dependent platelet and endothelial responses^17,18^.

The nonlinear flow response of this coupled valve–flow system remains unresolved. Previous valve-on-chip studies have primarily characterized changes in opening, recirculation, transport, and thrombotic response as functions of valve mechanics and imposed flow conditions. Whether post-valve flow undergoes a Reynolds-number-dependent symmetry-breaking transition, and how valve compliance or small bilateral geometric differences regulate its emergence, has not been established. Such a transition could amplify relatively modest structural or mechanical differences into substantially different downstream flow organizations.

The transport consequences of state selection are also important in blood, where the distribution of cellular and particulate components is shaped by the surrounding flow field. A persistent lateral asymmetry may therefore alter downstream delivery and clearance across the valve region. In the present study, RBC-scale particle transport was examined as a representative finite-size cellular transport process to determine whether flow-state selection produces persistent spatial redistribution.

Here, we tested the hypothesis that post-valve flow undergoes a Reynolds-number-dependent symmetry-breaking transition whose emergence is regulated by valve compliance and bilateral geometric bias. Using a biomimetic vein-on-a-chip platform, we combined computational fluid dynamics, fluid–structure interaction modeling, Ghost Particle Velocimetry (GPV), whole-blood velocimetry, and RBC-scale particle transport measurements to resolve the flow-state response across rigid and compliant valve configurations. We further examined whether transient forcing produces distinct increasing- and decreasing-Reynolds-number branches, whether the Reynolds-number-dependent asymmetric response remains detectable in whole blood, and whether lateral state selection produces persistent downstream transport heterogeneity.

## II. RESULTS

### A. Framework for resolving post-valve flow states

The venous-valve region was represented as a confined expansion comprising opposing leaflets, lateral sinus pockets, a restricted opening, and a downstream jet (Fig. 1A). This geometry permits qualitatively distinct organizations of the post-valve flow, including a centered, near-symmetric state and a laterally selected asymmetric state (Fig. 1B). Such symmetry breaking is well established in confined expansion flows and has been observed experimentally in geometrically symmetric systems^14^. We therefore examined whether increasing Reynolds number could trigger a comparable transition between post-valve flow states and how valve mechanics modified the emergence of asymmetry.

**FIG. 1.**
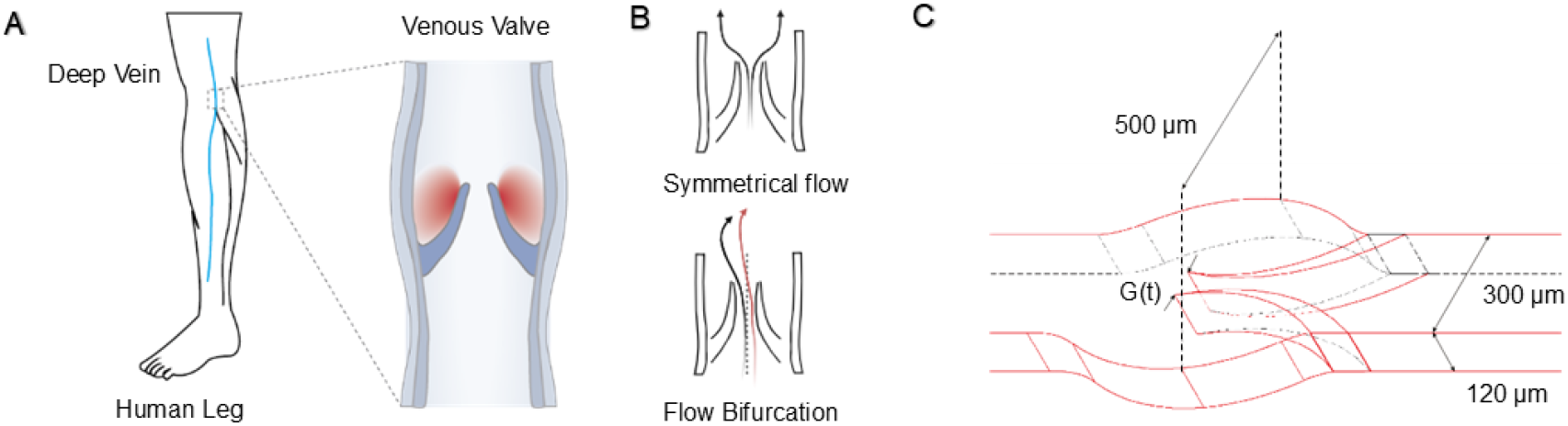
Conceptual framework and valve configurations for post-valve symmetry breaking. **(A)** Schematic of the venous-valve region, high-lighting the restricted leaflet opening, lateral valve pockets, and downstream expansion. **(B)** Conceptual post-valve flow states, showing a centered near-symmetric jet and a laterally selected asymmetric state. **(C)** Rigid fixed-gap and flexible-valve configurations used to distinguish intrinsic hydrodynamic symmetry breaking from flow–structure coupling. The rigid valve maintained a fixed opening, whereas the flexible-valve opening varied under fluid loading.

Rigid fixed-gap and flexible-valve configurations were used to distinguish the intrinsic hydrodynamic response from flow– structure coupling (Fig. 1C). The rigid configuration preserved the valve geometry under flow, whereas deformation of the flexible leaflets altered the effective opening and coupled valve mechanics to the evolving flow field. These configurations extend previous venous-valve-on-chip studies showing that leaflet mechanics influence valve opening, local hemodynamics, and thrombotic or particle localization^16,17^. Centered and geometrically biased flexible configurations were further used to distinguish the effects of compliance from those of bilateral geometric bias.

Post-valve flow organization was quantified using the mirror asymmetry index,

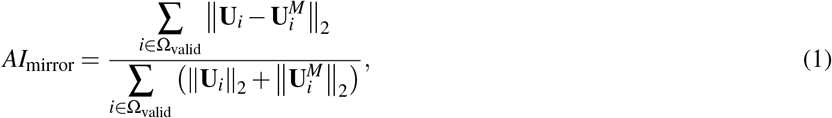

where **U**_*i*_ and 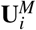 denote the velocity vector and its mirror-transformed counterpart at valid paired locations about the channel centerline, respectively. *AI*_mirror_ = 0 corresponds to ideal mirror symmetry, whereas larger values indicate progressively stronger bilateral flow asymmetry. Because the numerator is a vector norm, *AI*_mirror_ is intrinsically non-negative and quantifies asymmetry magnitude without encoding whether the selected state is displaced towards the left or right side of the channel.

Finite-size particle redistribution was quantified separately using the whole-field particle-event asymmetry index,

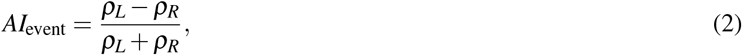

where *ρ*_*L*_ and *ρ*_*R*_ denote the area-normalized particle-event densities on the two sides of the channel. Downstream redistribution was additionally characterized using the section-based asymmetry index,

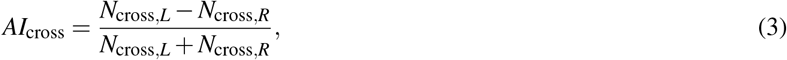

where *N*_cross,*L*_ and *N*_cross,*R*_ denote the cumulative numbers of section-associated particle-detection events on the two sides of the channel. Both particle indices retain the direction of lateral bias; where branch direction was not required, their absolute values were used to quantify the magnitude of transport asymmetry.

Together, these metrics provide complementary measures of continuous velocity-field organization and finite-size particle redistribution, allowing the hydrodynamic transition and its downstream transport consequences to be analyzed separately.

### B. Reynolds-number-dependent symmetry breaking and its regulation by valve compliance

In the geometrically symmetric rigid valve, increasing the instantaneous gap Reynolds number, *Re*_gap_, defined in Eq. (4), produced a marked reorganization of the post-valve flow (Fig. 2A). At low *Re*_gap_, the jet remained centered and the downstream field was approximately mirror symmetric. As *Re*_gap_ increased through the transition regime, the jet progressively lost bilateral symmetry and developed a lateral preference. At higher *Re*_gap_, the flow occupied a persistent asymmetric state characterized by a displaced jet and unequal organization of the two downstream regions.

**FIG. 2.**
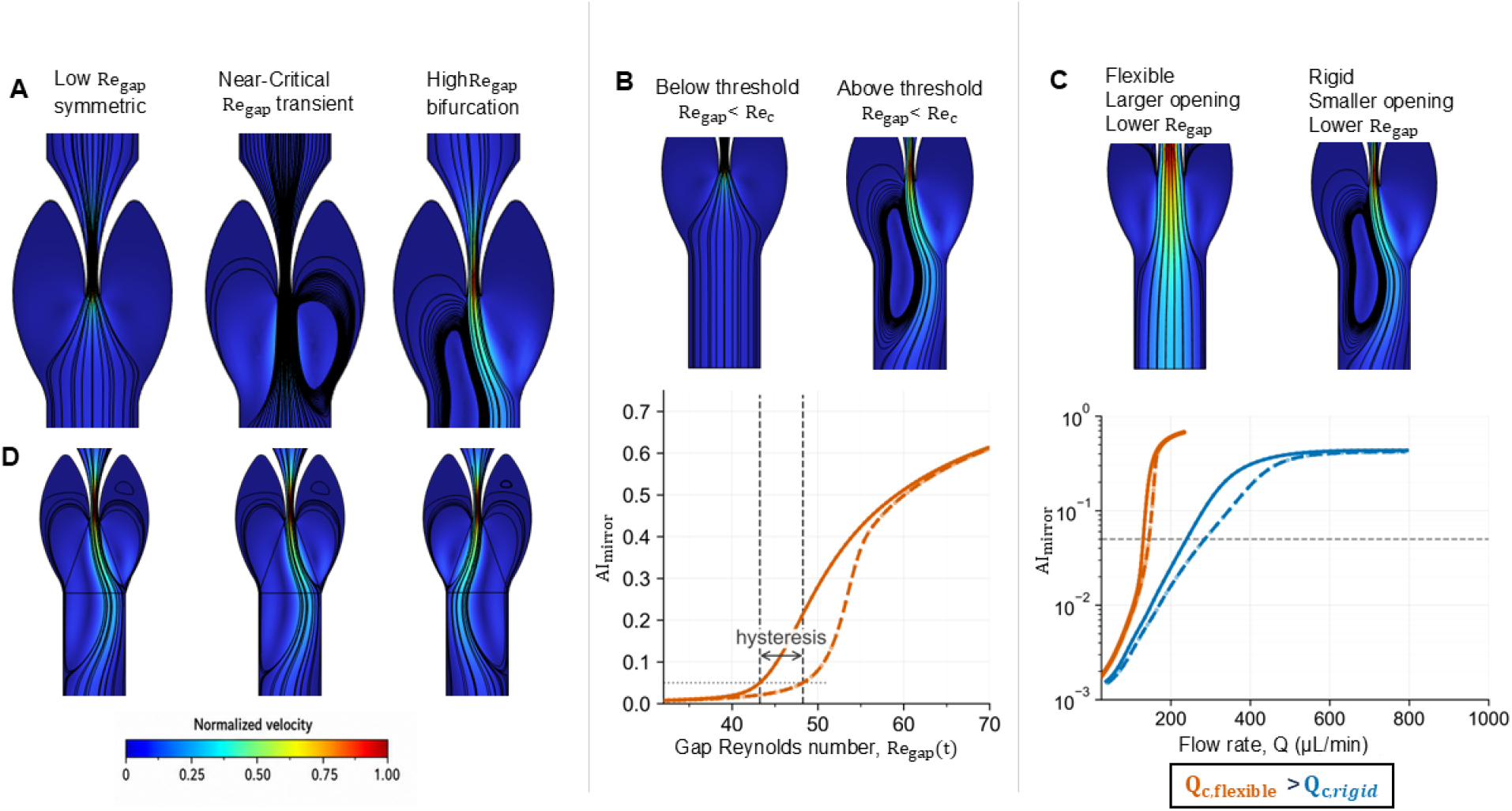
Reynolds-number-dependent symmetry breaking and its regulation by valve compliance. **(A)** Representative two-dimensional velocity fields for the geometrically symmetric rigid-valve model at low, near-transition, and high instantaneous *Re*_gap_, showing the progression from a centered near-symmetric jet through the transition regime to a symmetry-broken flow state. **(B)** Mirror asymmetry index, *AI*_mirror_, as a function of instantaneous *Re*_gap_ for the rigid fixed-gap valve. Solid and dashed curves denote the increasing- and decreasing-*Re*_gap_ branches, respectively. Each curve represents the median branch across the three simulated cycles. Their separation near the onset of pronounced asymmetry defines a reproducible hysteresis window under the applied transient forcing. **(C)** *AI*_mirror_ as a function of imposed flow rate, *Q*, for the rigid fixed-gap and centered flexible-valve models. Orange and blue curves denote the rigid and centered flexible valves, respectively, while solid and dashed curves denote increasing and decreasing *Q*. Flow-induced leaflet opening shifted the emergence of pronounced post-valve asymmetry to higher flow conditions in the centered flexible valve. **(D)** Post-valve velocity fields at peak forward flow for the three simulated cycles of the geometrically symmetric rigid-valve model, shown from left to right as cycles 1, 2, and 3. The jet selected the same lateral direction in the first two cycles and the opposite direction in the third, demonstrating access to mirror-related asymmetric flow states within the same symmetric geometry.

Because *AI*_mirror_ quantifies the magnitude of asymmetry without specifying its lateral direction, the peak forward-flow fields were also examined separately for each simulated cycle (Fig. 2D). The selected jet was deflected toward the same side during the first two cycles but switched to the opposite side during the third cycle. The emergence of opposite lateral states within the same geometrically symmetric model is consistent with selection between mirror-related asymmetric branches.

This transition was captured quantitatively by the mirror asymmetry index, *AI*_mirror_, defined in Eq. (1) (Fig. 2B). The index remained low in the near-symmetric regime before increasing markedly over a finite Reynolds-number range, distinguishing the emergence of the asymmetric state from a simple proportional increase in background asymmetry. The increasing- and decreasing-*Re*_gap_ phases followed distinct trajectories near the onset of pronounced asymmetry, defining a reproducible hysteresis window across the simulated cycles.

Valve compliance shifted this response (Fig. 2C). The centered flexible valve maintained lower *AI*_mirror_ over the low and intermediate flow range and developed pronounced asymmetry more gradually than the rigid fixed-gap valve. Flow-induced leaflet opening increased the effective gap and reduced the local throat velocity and associated inertial forcing, thereby extending the near-symmetric regime and shifting the emergence of pronounced lateral organization to higher imposed flow conditions.

The centered flexible valve nevertheless developed an asymmetric response at sufficiently high flow conditions. Compliance therefore delayed and broadened the emergence of symmetry breaking rather than eliminating the asymmetric flow state. These results identify flow-induced changes in valve opening as a mechanical control of the transition.

### C. GPV resolves the compliance-dependent flow response

GPV measurements provided experimental velocity fields for the rigid fixed-gap and centered flexible valves (Fig. 3A). At low *Re*_gap_, both configurations exhibited comparatively near-symmetric downstream flow. With increasing *Re*_gap_, the rigid valve developed a pronounced lateral displacement of the post-valve jet, whereas the centered flexible valve retained a more balanced flow distribution over a broader Reynolds-number range.

**FIG. 3.**
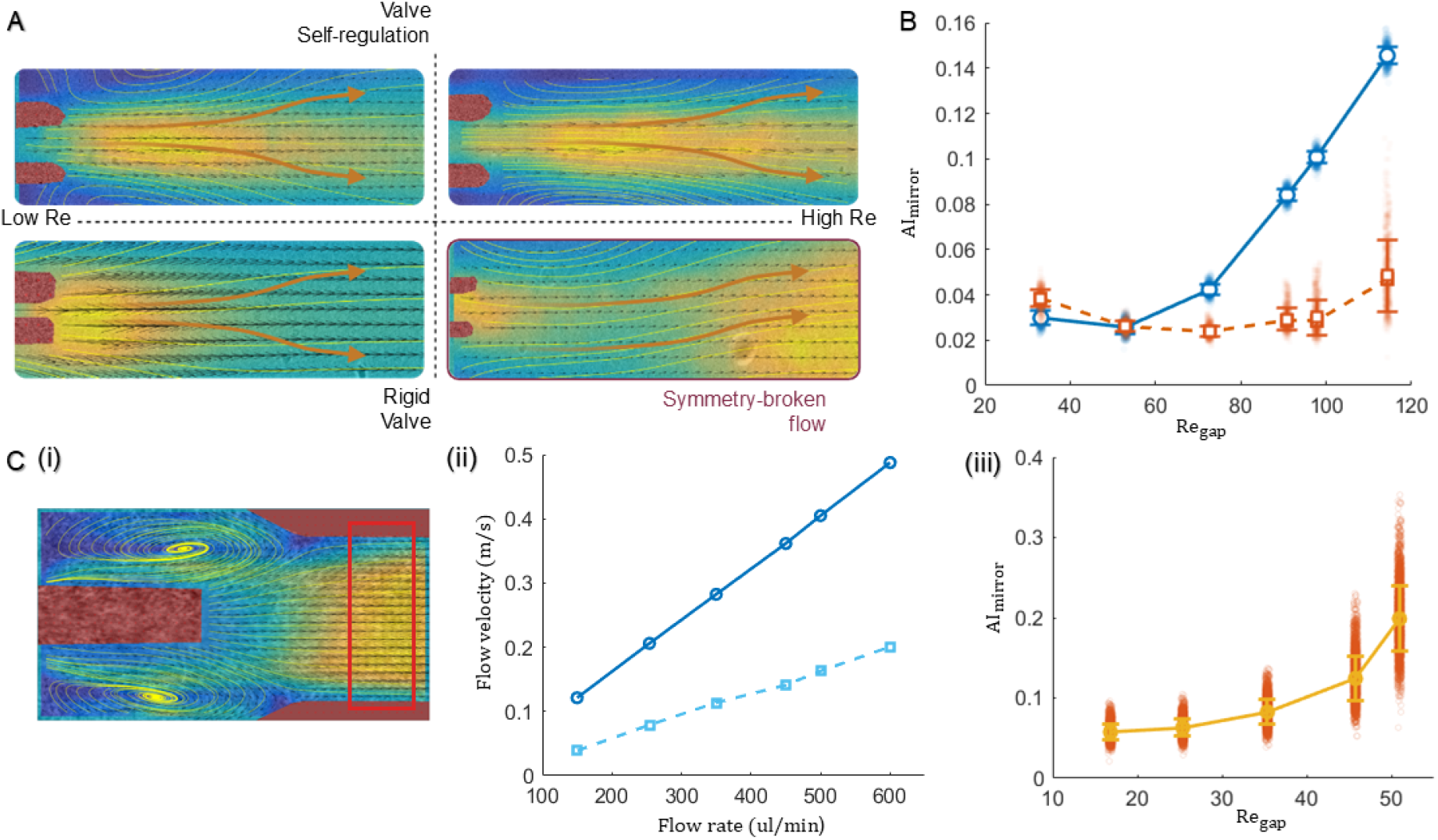
Experimental characterisation of post-valve flow using GPV and whole-blood imaging. **(A)** Representative velocity fields for the rigid and centered flexible valves at low and high *Re*_gap_. **(B)** *AI*_mirror_ as a function of *Re*_gap_ for the rigid and centered flexible valves, shown in blue and orange, respectively. **(C)** Whole-blood flow measurements. **(i)** Representative near-bottom velocity field. **(ii)** Relationship between velocities measured at the middle and near-bottom planes, shown in blue and orange, respectively. **(iii)** Whole-blood *AI*_mirror_ as a function of *Re*_gap_.

The corresponding mirror asymmetry index, *AI*_mirror_, defined in Eq. (1), showed the same compliance-dependent ordering (Fig. 3B). The rigid configuration exhibited a marked increase in velocity-field asymmetry near *Re*_gap_ *∼*50, while the centered flexible valve remained close to the near-symmetric state over the low and intermediate Reynolds-number range and developed pronounced asymmetry more gradually at higher *Re*_gap_.

The agreement in flow-state ordering between GPV and the computational results provides experimental support for the predicted effect of valve compliance. In both approaches, the centered flexible valve extended the near-symmetric regime relative to the rigid fixed-gap configuration. The experiment therefore shows that the compliance-dependent shift in post-valve asymmetry is a measurable feature of the physical system rather than a behavior confined to the numerical model.

### D. Whole blood retains the Reynolds-number-dependent asymmetry trend

Whole-blood measurements were used to assess whether the Reynolds-number-dependent asymmetry remained detectable under physiologically representative flow conditions. Because optical attenuation prevented reliable measurements at the channel mid-plane, velocimetry was performed at the near-bottom plane. An independent GPV calibration, obtained by measuring velocities at both the mid-plane and near-bottom plane under matched flow conditions, established the relationship between the two measurement planes (Fig. 3C(ii)). This relationship was subsequently used to convert the near-bottom whole-blood measurements to their corresponding mid-plane velocities.

Whole-blood velocity fields remained comparatively near-symmetric at low *Re*_gap_ and developed progressively stronger bilateral asymmetry as *Re*_gap_ increased (Fig. 3C). The response was more variable than that observed in the water-based GPV measurements, consistent with the cellular heterogeneity and shear-dependent rheology of whole blood^19,20^.

Despite this increased variability, the Reynolds-number-dependent ordering of the flow states was retained. The whole-blood measurements therefore show that the Reynolds-number-dependent asymmetric response remains detectable in a cellular, non-Newtonian fluid environment. This retained response supports the relevance of the observed flow-state organization under more physiologically representative flow conditions.

### E. Small geometric bias promotes an earlier asymmetric response in the flexible valve

To determine whether bilateral geometric bias modified the compliance-dependent delay in symmetry breaking, three valve configurations were compared under matched flow conditions: the rigid fixed-gap valve, the centered flexible valve, and a geometrically biased flexible valve (Fig. 4A).

**FIG. 4.**
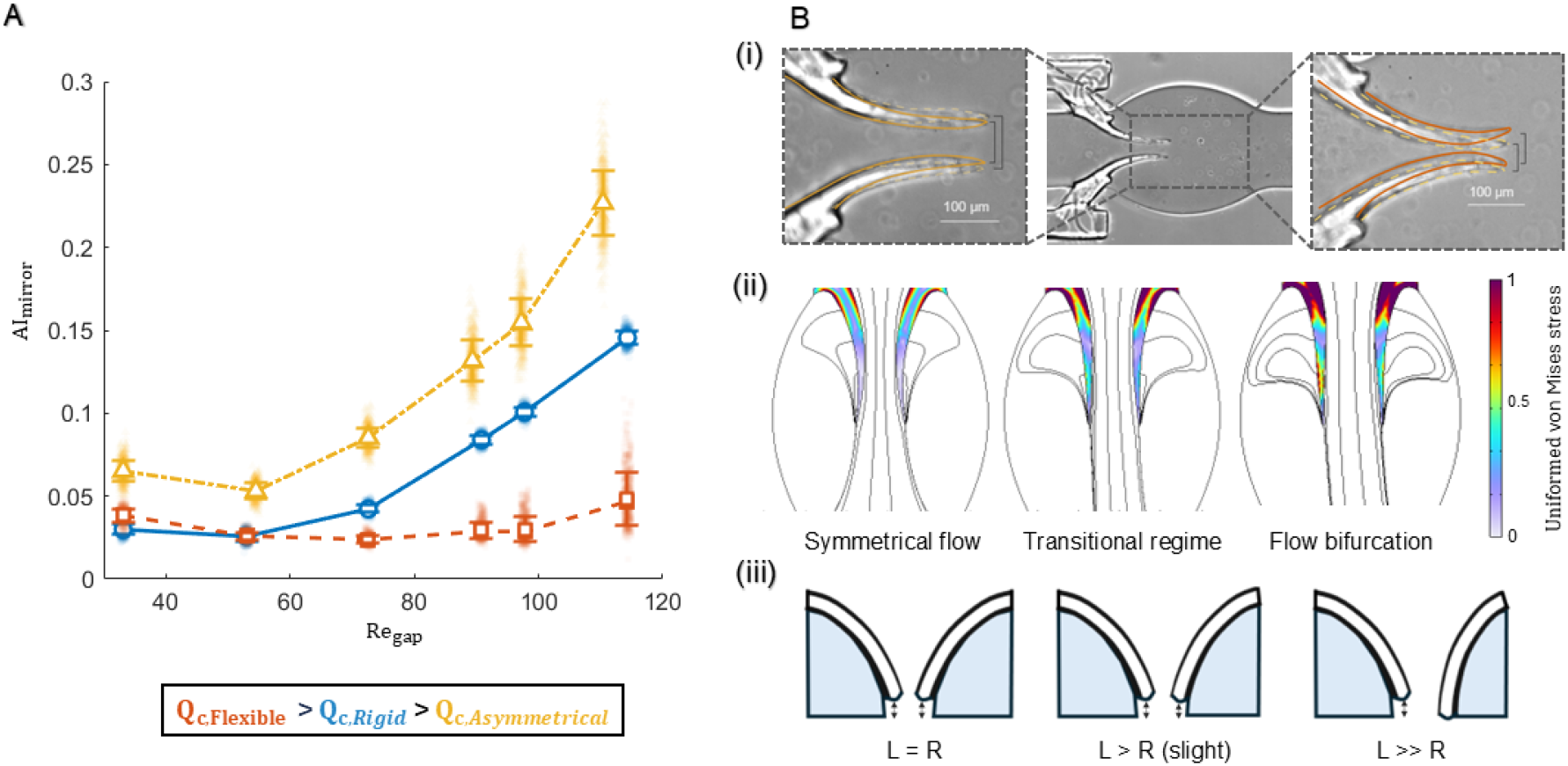
Effects of geometric bias and flow–structure interaction on flexible-valve behavior. **(A)** Reynolds-number-dependent *AI*_mirror_ for the rigid fixed-gap, centered flexible, and geometrically biased flexible valves. **(B)** Geometric and mechanical response of the flexible valves. **(i)** Experimentally measured valve geometries showing flow-induced opening and resting bilateral bias. Solid and dashed orange contours distinguish low- and high-*Re*_gap_ valve positions in the left image and geometrically biased and centered configurations in the right image. **(ii)** Fluid–structure interaction results showing the progressive development of asymmetric leaflet loading from the near-symmetric state through the transition regime to the symmetry-broken flow state. **(iii)** Schematic representation of the resulting bilateral differences in leaflet position and pocket geometry.

The centered flexible valve maintained the lowest *AI*_mirror_ over the low and intermediate Reynolds-number range. Introducing a small bilateral geometric bias substantially altered this response. Despite retaining compliant leaflets, the geometrically biased flexible valve developed pronounced asymmetry earlier and approached the sharper Reynolds-number-dependent response observed in the rigid configuration.

Microscopy showed that the imposed geometric bias was small relative to the channel dimensions (Fig. 4B(i)). The measured bilateral difference in tip-to-wall clearance was approximately 20 *µ*m, corresponding to approximately 6.7% of the channel width. Equivalently, this represented an opening-center displacement of approximately 10 *µ*m, or 3.3% of the channel width. Despite this modest perturbation, the Reynolds-number dependence of the post-valve flow changed markedly.

The centered and geometrically biased flexible valves showed only modest separation at low *Re*_gap_, but diverged strongly as the transition regime was approached. This Reynolds-number-dependent divergence is consistent with imperfection-sensitive branch selection in symmetry-breaking flows^11,12^. These results show that a small resting geometric bias can substantially promote the emergence of a laterally selected asymmetric state in the compliant valve.

### F. Flow asymmetry induces unequal leaflet deformation

Fully coupled fluid–structure interaction simulations were used to determine how the selected asymmetric flow state altered the mechanical response of the flexible valve (Fig. 4B(ii)). Before pronounced symmetry breaking, loading and deformation remained approximately symmetric between the two leaflets. As the flow progressed through the transition regime, the laterally selected state was accompanied by increasingly unequal mechanical loading of the two leaflets and a corresponding divergence in leaflet deformation.

At higher *Re*_gap_, the asymmetric mechanical response developed in concert with the displaced post-valve jet. Greater deformation on one leaflet and reduced deformation on the opposite leaflet altered the valve opening and pocket geometry (Fig. 4B(iii)), coupling the valve configuration to the selected flow branch. This behavior is consistent with a positive fluid–structure feedback mechanism in which the emerging asymmetric flow modifies the valve geometry in a manner that can mechanically reinforce the selected state.

The FSI results therefore show that the mechanical response of the flexible valve changes after lateral state selection, as asymmetric hydrodynamic loading drives unequal leaflet deformation associated with the selected flow organization.

### G. The symmetry-broken flow produces delayed and persistent RBC-scale transport asymmetry

RBC-scale polystyrene particles were used as finite-size probes to determine whether the post-valve flow transition produced a corresponding redistribution of the transported particulate phase. Particle transport was quantified from event-density maps accumulated over long image sequences without reconstructing complete particle trajectories (Fig. 5A). At low *Re*_gap_, particle events remained broadly distributed across the downstream channel, whereas increasing *Re*_gap_ produced a progressively lateralised distribution with a particle-rich pathway on one side and a persistent particle-depleted region on the other (Fig. 5B).

**FIG. 5.**
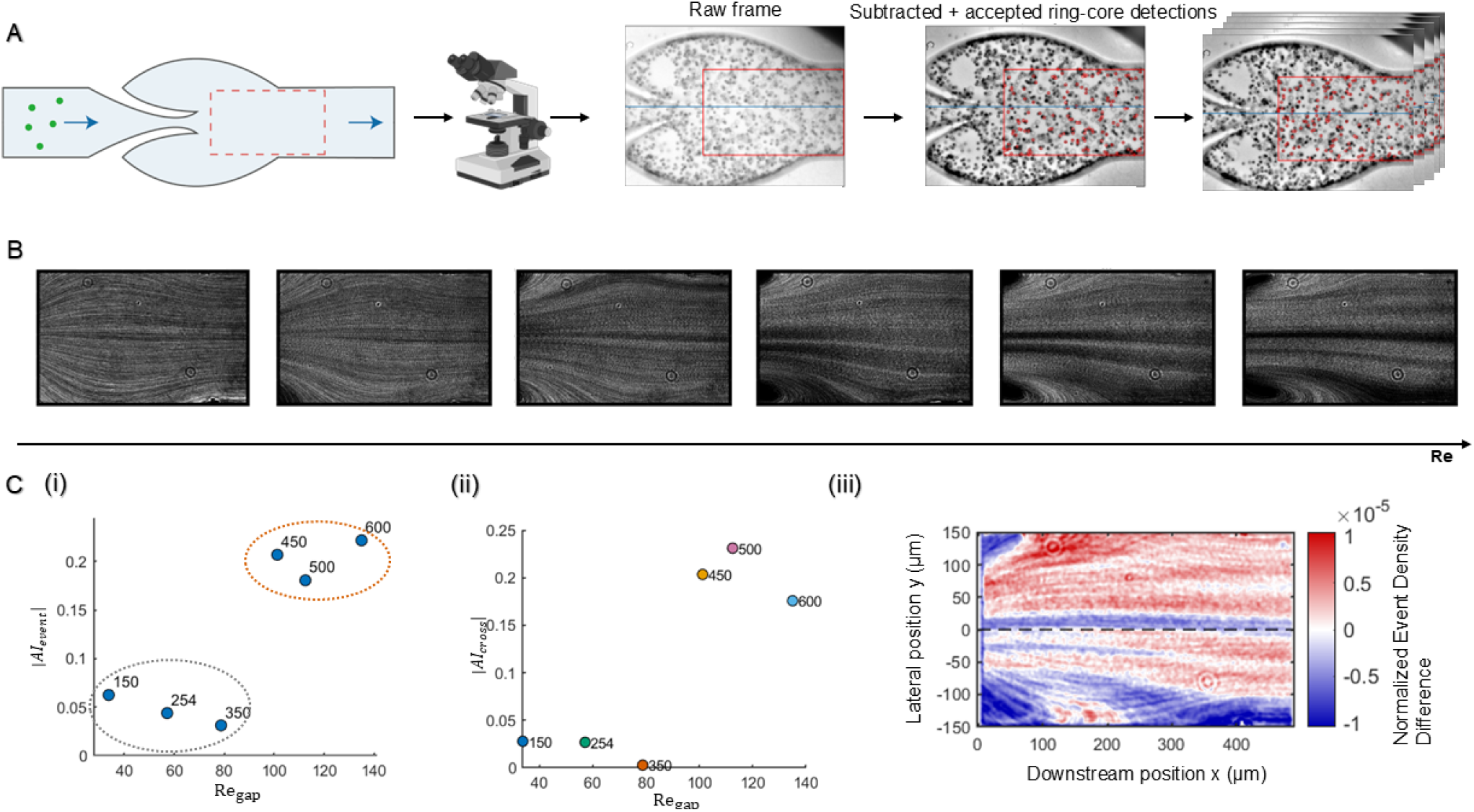
Reynolds-number-dependent redistribution of RBC-scale particles downstream of the rigid valve. **(A)** Particle-analysis workflow showing the downstream field of view, raw imaging, particle detection, and accumulated particle events. **(B)** Representative area-normalized particle-event density maps across increasing *Re*_gap_, showing the development of a laterally heterogeneous transport pattern at higher Reynolds number. **(C)** Quantification and spatial characterisation of particle redistribution. **(i)** Absolute whole-field particle-event asymmetry, |*AI*_event_|, as a function of *Re*_gap_. **(ii)** Absolute downstream section-based asymmetry, |*AI*_cross_|, as a function of *Re*_gap_. **(iii)** Difference between the highest- and lowest-*Re*_gap_ particle-event density fields. Positive and negative values indicate regions of relative particle enrichment and depletion at high Reynolds number, respectively; the dashed line marks the channel centerline.

The magnitude of this redistribution was quantified using the whole-field particle-event asymmetry index, *AI*_event_, defined in Eq. (2). Lower-Reynolds-number measurements remained at comparatively low *AI*_event_, whereas a marked increase emerged only at higher *Re*_gap_ (Fig. 5C(i)). This increase occurred after velocity-field asymmetry had already become detectable, indicating that the finite-size particle distribution responded later than the continuous velocity field. This separation indicates that pronounced particle redistribution emerged at higher *Re*_gap_ than the detectable velocity-field asymmetry.

Persistence of the redistribution was assessed using the downstream section-based asymmetry index, *AI*_cross_, defined in Eq. (3). Elevated |*AI*_cross_| at downstream sampling locations showed that the lateral particle bias persisted beyond the immediate post-valve region after the particles entered the straight downstream channel (Fig. 5C(ii)). The spatial structure of this redistribution was further resolved by subtracting the lowest-*Re*_gap_ particle-event density field from the highest-*Re*_gap_ field (Fig. 5C(iii)). The resulting difference map showed spatially separated regions of relative particle enrichment and depletion at high Reynolds number, demonstrating a lateral reorganization of downstream particle transport.

The symmetry-breaking flow transition was therefore followed by a delayed and persistent redistribution of RBC-scale particles.

## III. DISCUSSION

### A. A symmetry-breaking transition in post-valve venous flow

The principal finding of this study is a Reynolds-number-dependent symmetry-breaking transition in post-valve venous flow. A centered near-symmetric jet evolved into a persistent asymmetric state as the local inertial forcing increased, including in the geometrically symmetric rigid-valve configuration. Pronounced lateral organization can therefore emerge from the post-valve flow dynamics without a substantial pre-existing geometric bias.

This behavior is closely related to the symmetry-breaking bifurcations reported in symmetric sudden expansions and other confined expansion flows, where increasing inertial forcing destabilizes a centered laminar jet and promotes mirror-related asymmetric states^11–14^. Consistent with this behavior, the symmetric rigid-valve model accessed both mirror-related lateral states across successive forcing cycles, indicating that the direction of post-valve deflection is itself a selectable feature of the symmetry-broken flow state. The valve geometry introduces additional features absent from canonical expansion flows, including opposing leaflets, lateral pockets, and an opening that can change under fluid loading. These features make the transition sensitive to both local geometry and structural mechanics^15–18^.

The symmetry-breaking response adds a flow-state description to conventional hemodynamic quantities such as velocity, shear stress, recirculation, and residence time^3,4,8^. Those quantities remain essential for characterizing local hemodynamics, while the present results show that the spatial organization of the post-valve flow can itself change qualitatively. Near the transition regime, relatively small changes in local inertial conditions can therefore reorganize the downstream flow pattern as a whole.

Reynolds number remained the primary control parameter governing this transition. The transient simulations additionally revealed repeatable separation between the increasing- and decreasing-*Re*_gap_ branches, consistent with hysteresis near the onset of pronounced asymmetry. This branch separation was reproducible across successive flow cycles. Its detailed dynamical origin and dependence on the forcing trajectory remain to be resolved.

The transition becomes particularly important once the selected flow organization is transferred to finite-size transport. In that regime, lateral asymmetry in the velocity field produces persistent differences in downstream particle distribution and therefore alters the spatial structure of delivery and clearance around the valve.

### B. Valve mechanics regulate the emergence of asymmetry

Valve compliance exerted a state-dependent influence on post-valve flow organization. Before lateral state selection, approximately symmetric leaflet opening increased the effective valve gap, reduced the local throat velocity and associated inertial forcing, and extended the near-symmetric regime. The centered flexible valve consequently developed pronounced asymmetry at higher flow conditions and over a broader transition range than the rigid fixed-gap configuration. Previous vein-on-chip studies have shown that leaflet mechanics alter valve opening, local hemodynamics, and the spatial localization of particles or thrombotic material^16,17^. The present results show that compliance also regulates the local inertial conditions governing the emergence of symmetry breaking.

The instantaneous *Re*_gap_ captures both the local gap velocity and the evolving valve opening and therefore provides a direct measure of the inertial conditions within the constriction. In the centered flexible valve, flow-induced opening reduced the acceleration through the valve gap for a given imposed flow condition, shifting the development of pronounced asymmetry to higher flow conditions.

The mechanical role of compliance changed after lateral state selection. At lower flow conditions, the two leaflets in the symmetric fluid–structure interaction model experienced similar loading and deformation. Once the post-valve jet became asymmetric, hydrodynamic loading diverged between the two sides and produced unequal leaflet deformation despite identical material properties and symmetric boundary conditions. The resulting changes in leaflet position modified the local opening and pocket geometry in a direction consistent with mechanical reinforcement of the selected flow organization.

Geometric bias provided a second route to lateral state selection. The geometrically biased flexible valve differed from the centered configuration by a small resting perturbation: the measured difference in tip-to-wall clearance was approximately 20 *µ*m, corresponding to 6.7% of the channel width, with an opening-center displacement of approximately 10 *µ*m, corresponding to 3.3% of the channel width. The difference in flow response remained modest at low Reynolds number and increased markedly as the transition regime was approached. This behavior is consistent with imperfection-sensitive symmetry breaking in confined expansion flows, where small perturbations become increasingly influential near the loss of the symmetric flow organization^11–14^.

The combined results indicate that valve mechanics regulate several stages of the transition. Symmetric opening shifts the conditions under which pronounced asymmetry emerges, geometric bias promotes lateral state selection, and asymmetric deformation can reinforce the selected organization after transition. Anatomical variability in valve geometry and leaflet mechanics may therefore generate substantial differences in post-valve flow organization under otherwise similar imposed flow conditions^15^.

### C. From flow asymmetry to persistent RBC-scale transport heterogeneity

The symmetry-breaking transition also reorganized the transport of RBC-scale objects. GPV resolved the velocity-field response, while the 8 *µ*m polystyrene particles quantified downstream redistribution at an RBC-relevant length scale. *AI*_mirror_ therefore characterizes the emergence of velocity-field asymmetry, whereas *AI*_event_ and *AI*_cross_ quantify its transport consequences.

The velocity-field response preceded pronounced lateral particle redistribution. This delay is consistent with the accumulation of transport differences as particles move through the selected asymmetric flow field over time and downstream distance. Once established, the particle asymmetry persisted into the downstream straight channel, extending the influence of state selection beyond the immediate post-valve jet.

The spatial measurements showed that symmetry breaking reorganized particle transport into laterally distinct pathways. At higher *Re*_gap_, one side of the downstream channel became relatively enriched in particle events while the opposite side became depleted, as resolved by the high-minus-low-*Re*_gap_ event-density difference map. Elevated downstream |*AI*_cross_| further showed that this organization persisted after the particles left the valve region. The selected flow state therefore generated a spatially persistent pattern of RBC-scale transport.

This transport heterogeneity creates unequal delivery and clearance conditions across the valve region. A dominant pathway preferentially conveys blood-borne material through one side of the downstream channel, producing spatially distinct exposure conditions that are absent in the centered flow state. Such differences provide a fluid-mechanical basis for heterogeneity in washout, cellular delivery, and local hemodynamic exposure relevant to venous thrombosis^3,4,8^.

The whole-blood measurements extend this picture to a cellular, shear-dependent fluid. The Reynolds-number-dependent asymmetric flow organization remained detectable under these conditions^19,20^, indicating that the flow-state response persists in a substantially more complex rheological environment. Together, the velocity-field, whole-blood, and particle measurements connect post-valve symmetry breaking with persistent heterogeneity in downstream transport.

### D. Limitations and future work

The computational models were structured to separate the hydrodynamic transition from the effects of valve deformation. The rigid fixed-gap configuration isolated the Reynolds-number-dependent flow response, while the fluid–structure interaction model introduced compliant opening and asymmetric hydrodynamic loading. The latter used idealized leaflet properties, attachment conditions, and reduced geometric complexity. Three-dimensional leaflet geometry, mechanical heterogeneity, and coupling to the vessel wall may shift the quantitative transition range and the magnitude of fluid–structure coupling in native valves.

The instantaneous gap Reynolds number, *Re*_gap_, provided a local measure of the inertial conditions within the evolving valve opening. The present parameter sampling resolved a finite transition regime together with repeatable separation of the increasing- and decreasing-*Re*_gap_ branches. A formal reconstruction of the transition structure will require denser Reynolds-number sampling, controlled upward and downward forcing protocols, longer recordings, and targeted perturbations. These studies could determine the dynamical origin of the observed hysteresis and distinguish the contributions of transient relaxation, forcing rate, and prior flow evolution.

Whole-blood velocimetry was restricted to the near-bottom plane and converted to the corresponding mid-plane velocity using the independent GPV calibration. Because blood viscosity varies with shear rate, hematocrit, deformability, aggregation, temperature, and confinement^19,20^, the quantitative mapping between imposed flow rate, local velocity, and effective *Re*_gap_ carries additional uncertainty. The Reynolds-number-dependent ordering of the asymmetric response nevertheless remained detectable in whole blood. Direct rheological characterisation and three-dimensional velocimetry would allow the influence of non-Newtonian effects on the quantitative transition behavior to be resolved more fully^21^.

The 8 *µ*m polystyrene particles capture an RBC-relevant transport scale with simpler mechanical properties than native erythrocytes. Deformability, cell–cell interactions, and migration may alter the quantitative redistribution observed in blood, while the whole-field event-based analysis quantifies accumulated detections, whereas the section-based analysis reconstructs short trajectories to identify individual crossing events. Future measurements using deformable red cells, platelets, and endothelialized valve models could directly resolve branch-dependent cellular delivery, wall encounter, clearance, and thrombotic deposition. A further extension is to couple thrombus growth with flow-state selection, allowing evolving changes in valve geometry and local resistance to modify the subsequent hemodynamics and transport.

## IV. METHODS

### A. Valve-on-a-chip fabrication and valve configurations

Microfluidic devices were fabricated using the valve-on-a-chip procedure previously established for compliant venous-valve microfluidic models^16,17^. Briefly, SU-8 2075 photoresist was patterned on a silicon wafer by conventional photolithography to produce a negative-relief mould. Poly(dimethylsiloxane) (PDMS; Sylgard 184 Silicone Elastomer Kit, Dow Corning) was mixed with curing agent at a 10:1 mass ratio, degassed under vacuum, cast over the mould, and cured at 70 ^*°*^C for 2 h. Inlet and outlet ports were formed using a 1.5-mm biopsy punch, and the moulded PDMS layer was bonded to a PDMS-coated glass substrate following corona-discharge activation.

Flexible valve leaflets were fabricated in situ by projection photopolymerisation using the formulation reported previously^16^. The precursor contained 50% (w/v) poly(ethylene glycol) diacrylate (PEGDA; *M*_r_ *≈*575, Sigma–Aldrich) and 4% (w/v) 2-hydroxy-2-methylpropiophenone photoinitiator (Sigma–Aldrich). The leaflet mask was projected through a 20*×* objective, and the precursor was crosslinked using approximately 365-nm ultraviolet illumination for 300 ms. The channels were subsequently flushed with deionised water to remove uncrosslinked precursor.

The devices contained a 300 *µ*m-wide, 120 *µ*m-high rectangular channel with two opposing valve leaflets, adjacent lateral sinus pockets, and a downstream expansion. Three valve configurations were examined: a rigid fixed-gap valve, a centered flexible valve, and a geometrically biased flexible valve. The rigid valve had a minimum opening of approximately 46 *µ*m and preserved a fixed constriction geometry during flow. Both flexible configurations used photopolymerised PEGDA leaflets within the same channel architecture. The centered configuration was designed with approximately matched leaflet positions, whereas the geometrically biased configuration incorporated a small bilateral difference in resting leaflet position. Valve opening and bilateral geometric bias were quantified from calibrated bright-field images as described in Section IV G.

### B. Flow conditions and gap-based Reynolds number

Experimental flow was generated using a Legato 111 programmable syringe pump (KD Scientific, Holliston, MA, USA). Each prescribed flow rate was imposed as a separate steady condition and maintained for 5 min before image acquisition. The imposed flow rate remained constant throughout the corresponding recording. No continuous flow-rate ramp or oscillatory waveform was applied during the experimental measurements.

Water-based measurements with the rigid fixed-gap valve, centered flexible valve, geometrically biased flexible valve, and 8 *µ*m particle suspension were performed at imposed volumetric flow rates of 150, 254, 350, 450, 500, and 600 *µ*Lmin^*−*1^. Whole-blood measurements were performed at 600, 889, 1225, 1575, and 1750 *µ*Lmin^*−*1^ to access a comparable range of local inertial conditions. Transient bidirectional forcing was used only in the computational models described in Sections IV C and IV D.

Because the strongest local acceleration occurred through the valve opening and the subsequent expansion governed the downstream flow reorganization^11–13^, Reynolds number was defined using the local gap velocity and the corresponding valve opening,

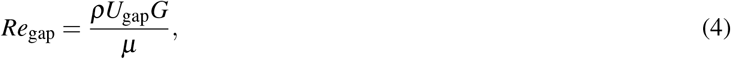

where *ρ* is the fluid density, *µ* is the dynamic viscosity, *U*_gap_ is the velocity magnitude evaluated at the defined gap sampling location, and *G* is the corresponding valve-gap width.

For the rigid fixed-gap valve, *G ≈*46 *µ*m. In the water-based experiments, *U*_gap_ was obtained from the measured middle-plane velocity field at the center of the valve opening. For the flexible-valve experiments, the valve opening was measured independently after the 5-min equilibration period at each imposed flow condition, and the corresponding measured value of *G* was used in Eq. (4). In the computational models, *U*_gap_(*t*) was evaluated at a fixed sampling location positioned at the center of the resting valve opening; for the flexible model, the instantaneous opening *G*(*t*) was obtained separately from the deformed leaflet geometry.

For the water-based experimental measurements, *ρ*_water_ = 997 kgm^*−*3^ and *µ*_water_ = 8.9 *×* 10^*−*4^ Pas were used. Whole-blood Reynolds-number scaling used *ρ*_blood_ = 1060 kgm^*−*3^ and *µ*_blood_ = 3.115 *×* 10^*−*3^ Pas. The fluid properties used in the computational models are specified separately in Sections IV C and IV D.

Whole-blood velocity fields were acquired approximately 10 *µ*m above the lower channel surface because optical attenuation prevented reliable measurements at the channel mid-plane. An independent water-based two-plane calibration was therefore used to relate the near-bottom velocity scale to the corresponding middle-plane velocity scale before evaluating Eq. (4). The calibration procedure is described in Section IV F.

### C. Rigid-valve computational model

Time-dependent computational fluid dynamics simulations were performed in COMSOL Multiphysics 6.2 (COMSOL AB, Stockholm, Sweden) using the *Laminar Flow* interface. The two-dimensional computational domain represented the central longitudinal plane of the rigid fixed-gap valve described in Section IV A, preserving the principal constriction–expansion geometry characteristic of symmetry-breaking internal flows^11–13^. The fluid was treated as incompressible and Newtonian, with conservation of mass and momentum governed by

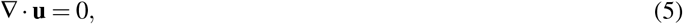

And

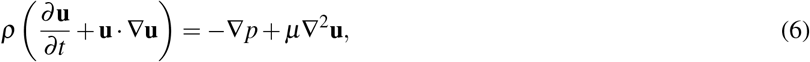

where **u** is the velocity vector, *p* is pressure, *ρ* is fluid density, and *µ* is dynamic viscosity. The baseline dynamic viscosity was set to *µ*_0_ = 1.0 *×*10^*−*3^ Pas, and the fluid density was obtained from the built-in COMSOL liquid-water material model evaluated at 293.15 K. No-slip conditions were applied to the channel walls and rigid valve surfaces.

The model was driven by a periodic pressure difference corresponding to a 3-s bidirectional flow cycle. Each cycle comprised a 2-s phase in one axial direction followed by a 1-s phase in the opposite direction, with a smooth squared-sine temporal profile within each phase. The imposed flow therefore returned to zero at the beginning of the cycle, at flow reversal, and at the end of the cycle. The temporal waveform was shared across the computational configurations, while the pressure amplitude was calibrated separately for each geometry to obtain the prescribed velocity range. A numerical viscosity-penalization treatment, based on the use of locally increased viscosity to suppress flow within penalised regions^22^, was also included as part of the computational implementation. Additional details of the waveform, pressure calibration, and viscosity-penalization implementation are provided in the Supplementary Methods.

The rigid-valve model was solved from 0 to 9 s using a time-dependent formulation, corresponding to three complete forcing cycles, with solution fields stored at 5-ms intervals. The domain was discretized using an unstructured finite-element mesh with local refinement near the leaflet tips, within the valve opening, and in the adjacent downstream valve region. At each stored time point, the local gap velocity, *U*_gap_(*t*), was obtained from the COMSOL velocity magnitude (spf.U) evaluated at a fixed sampling point positioned midway between the two rigid leaflet tips. The instantaneous gap-based Reynolds number was then calculated using Eq. (4) with *G≈* 46 *µ*m and the baseline viscosity *µ*_0_.

Post-valve velocity-field asymmetry was quantified using *AI*_mirror_ defined in Eq. (1). Each of the three forcing cycles was analyzed independently. Within the forward-flow phase of each cycle, the increasing- and decreasing-*Re*_gap_ branches were identified from the temporal evolution of *Re*_gap_ and retained separately. For the *Re*_gap_–*AI*_mirror_ comparison, the cycle-resolved branches were linearly interpolated onto a common *Re*_gap_ grid, and the median *AI*_mirror_ across the three cycles was calculated at each grid point to obtain representative increasing and decreasing branches. Peak-forward-flow velocity fields were also extracted separately from each cycle to resolve the lateral direction of the selected asymmetric state. Velocity fields, *U*_gap_, instantaneous *Re*_gap_, phase and cycle information, and *AI*_mirror_ were exported for subsequent analysis in MATLAB.

### D. Flexible-valve fluid–structure interaction model

The flexible-valve model retained the fluid formulation, transient forcing, temporal sampling, and post-processing framework described for the rigid-valve model in Section IV C, with the rigid leaflets replaced by deformable PEGDA structures. Fluid– structure interaction simulations were performed in COMSOL Multiphysics 6.2 by coupling the *Laminar Flow, Solid Mechanics*, and *Moving Mesh* interfaces through the *Fluid–Structure Interaction* multiphysics coupling. The two leaflets were assigned identical initial geometry, material properties, attachment conditions, and resting positions. Their roots were fixed, while the remaining leaflet boundaries were free to deform under fluid loading. Fluid traction was transferred to the leaflet surfaces, and the resulting structural displacement was returned to the fluid domain through the moving mesh, with no slip imposed at the deforming fluid–solid interfaces.

The PEGDA leaflets were represented as nearly incompressible hyperelastic solids using a two-parameter Mooney–Rivlin formulation. Structural motion was governed by

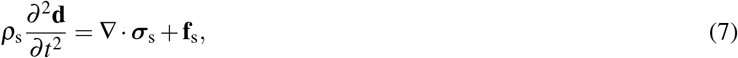

where *ρ*_s_ is the solid density, **d** is the displacement vector, ***σ***_s_ is the solid stress tensor, and **f**_s_ is the body-force density. The assigned leaflet density was *ρ*_s_ = 1100 kgm^*−*3^. The implemented Mooney–Rivlin coefficients were *C*_10_ = 23.492 kPa and *C*_01_ = 2.3492 kPa, with a bulk modulus of *K* = 1.0 *×* 10^7^ Pa. These coefficients correspond to an initial shear modulus of approximately 51.7 kPa and, in the nearly incompressible small-strain limit, an equivalent Young’s modulus of approximately 155 kPa. The same constitutive properties were applied to both leaflets.

Leaflet deformation was quantified from the displacement of each free tip,

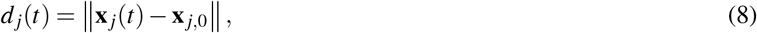

where **x** _*j*,0_ and **x** _*j*_(*t*) are the resting and deformed tip positions of leaflet *j*, respectively. The instantaneous valve opening was defined from the transverse separation of the two opposing leaflet tips,

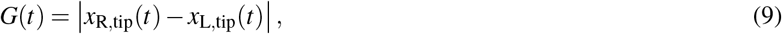

and bilateral deformation asymmetry was quantified as

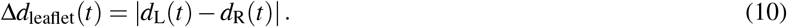

The local gap velocity, *U*_gap_(*t*), was evaluated from spf.U at the same fixed sampling location defined at the center of the resting valve opening; this point did not move with leaflet deformation. The instantaneous *Re*_gap_ was therefore calculated using Eq. (4) with the simulated opening *G*(*t*) and the baseline fluid viscosity. The flexible geometry used its own calibrated pressure amplitude and viscosity-penalization implementation within the common computational forcing framework. For the imposed-flow-rate comparison between rigid and flexible models, increasing- and decreasing-*Q* branches were treated separately. The cycle-resolved data were interpolated on a common linear *Q* grid in log_10_(*AI*_mirror_) space, and the median across repeated cycles was calculated in the same space before conversion back to *AI*_mirror_ for presentation. Additional constitutive-model details, geometry-specific pressure calibration, and the viscosity-penalization treatment are provided in the Supplementary Methods.

### E. Ghost-particle velocimetry and velocity-field analysis

Water-based velocity fields were measured using ghost-particle velocimetry (GPV), following the optical and image-analysis principles established previously^23,24^. The working fluid consisted of deionised water containing 200-nm polystyrene particles (Sigma–Aldrich, Merck) at a concentration of 0.1% and Triton X-100 at 0.01%. The nanoparticles were below the optical diffraction limit and generated a dynamic speckle pattern under partially coherent bright-field illumination. Imaging was performed using a Nikon Eclipse Ti2 inverted microscope (Nikon, Japan) equipped with a 20 *×*objective and a Photron FASTCAM Nova-series high-speed camera (Photron, Japan). The condenser aperture was adjusted to provide an illumination numerical aperture of approximately 0.15–0.20, consistent with previously reported GPV implementations^23,24^. For the water-based measurements, the focal plane was positioned near the channel mid-plane.

Each prescribed flow rate was imposed as a separate steady condition using the protocol described in Section IV B. Image sequences were acquired at a high frame rate selected according to the local flow velocity to maintain suitable inter-frame displacement of the GPV speckle pattern. The exposure time was fixed at 4 *µ*s. Static optical features were removed by temporal-median background subtraction before velocity-field reconstruction.

The GPV image sequences were analyzed in MATLAB using PIVlab version 3.06^25^. Consecutive image pairs were evaluated using fast-Fourier-transform window-deformation cross-correlation with linear window deformation. A two-pass interrogation scheme was used, with 32 *×*32-pixel and 16 *×*16-pixel interrogation windows in the first and second passes, respectively, and a step size equal to one-half of the corresponding interrogation-window dimension. The resulting two-dimensional velocity fields were exported as frame-resolved spatial and velocity arrays for subsequent analysis.

Post-valve velocity-field asymmetry was quantified using *AI*_mirror_ defined in Eq. (1). The geometric channel centerline was determined independently from calibrated bright-field images by fitting the two straight channel walls, and the velocity field was transformed into a channel-aligned coordinate system before mirror comparison. Reflection preserved the along-channel velocity component and reversed the channel-normal component. Only spatial locations containing valid velocity vectors in both the original and mirror-transformed fields were included in the calculation. The same centerline definition and post-valve analysis procedure were applied across all directly compared conditions.

For each recording, *AI*_mirror_ was calculated independently for each velocity field before condition-level summarisation. The local gap velocity *U*_gap_ used in Eq. (4) was obtained from the measured velocity magnitude at the center of the valve opening. For the rigid fixed-gap valve, the fixed opening was used, whereas the flow-specific measured opening was used for the flexible-valve experiments. Detailed coordinate transformation, interpolation, masking, vector-selection, quality-control, and frame-level filtering procedures are provided in the Supplementary Methods.

### F. Whole-blood velocimetry and velocity-plane calibration

Non-identifiable, citrate-anticoagulated human whole-blood samples were obtained from the Australian Red Cross Lifeblood Biological Resources program. The samples had been collected with donor consent for research purposes and remained citrate-anticoagulated throughout the velocimetry experiments. Samples were stored at 4 ^*°*^C and used within 7 days. Before perfusion, the blood was warmed to 37 ^*°*^C and mixed gently to restore cellular homogeneity. Whole-blood measurements were performed in the rigid fixed-gap valve under the steady-flow conditions described in Section IV B.

Owing to the optical attenuation and scattering produced by whole blood, reliable velocity measurements could not be obtained at the channel mid-plane. The focal plane was therefore positioned approximately 10 *µ*m above the lower channel surface. Imaging was performed using the optical and high-speed imaging system described in Section IV E. The acquisition frame rate was selected according to the local flow velocity, and the exposure time was fixed at 4 *µ*s. Static optical features were removed by temporal-median background subtraction.

The moving image texture generated by the cellular components of whole blood was analyzed in MATLAB using PIVlab version 3.06^25^, using the same FFT-based window-deformation cross-correlation procedure described for the water-based velocity measurements in Section IV E. The resulting two-dimensional near-bottom velocity fields were exported as frame-resolved spatial and velocity arrays. Post-valve asymmetry was quantified using *AI*_mirror_ defined in Eq. (1), with the same geometry-based centerline definition and mirror-analysis procedure used for the water-based measurements.

An independent water-based two-plane calibration was used to relate the near-bottom velocity scale accessible in whole blood to the corresponding middle-plane velocity scale. Water-based GPV measurements were acquired at both the near-bottom and middle planes over matched imposed flow conditions, and the relationships between imposed flow rate and local velocity were fitted independently for the two planes. The near-bottom velocity associated with each whole-blood condition was mapped through these calibration relationships to the corresponding middle-plane velocity scale. This calibrated velocity was then used as *U*_gap_ in the gap-based Reynolds-number definition given by Eq. (4), together with the rigid valve opening and the blood properties specified in Section IV B. The complete two-plane calibration equations and fitted coefficients are provided in the Supplementary Methods.

For each recording, *AI*_mirror_ was calculated independently from the frame-resolved velocity fields before condition-level summarisation. Detailed interpolation, vector-selection, quality-control, frame-exclusion, and statistical procedures were identical to those applied to the water-based velocity fields and are provided in the Supplementary Methods. Whole blood exhibits shear-dependent cellular rheology^19,20^; the effective viscosity adopted in Eq. (4) was used as the common Reynolds-number scaling for comparison across the tested whole-blood conditions.

### G. Flexible-valve geometry and geometric-bias quantification

Flexible-valve geometry was quantified from calibrated bright-field images to measure the valve opening and bilateral leaflet position under steady flow. Images were acquired under the no-flow condition and after equilibration at each imposed flow rate. The positions of the two opposing leaflet tips and the adjacent channel walls were identified from the original microscopy images using the same geometric landmarks across all conditions.

The valve opening was defined as the transverse separation between the two leaflet tips,

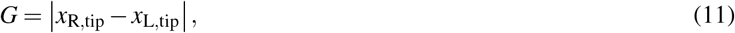

where *x*_L,tip_ and *x*_R,tip_ are the transverse coordinates of the left and right leaflet tips, respectively. The flow-induced change in opening was calculated relative to the no-flow configuration as

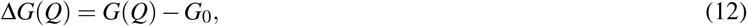

where *G*_0_ is the resting opening and *G*(*Q*) is the opening measured at imposed flow rate *Q*. The measured opening at each condition was used in the corresponding gap-based Reynolds-number calculation defined in Eq. (4).

Bilateral leaflet position was characterized using the transverse clearance between each leaflet tip and its adjacent channel wall,

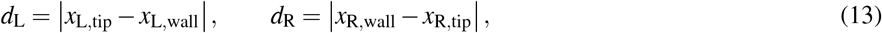

and the magnitude of geometric bias was defined as

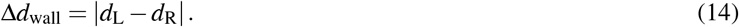

For approximately parallel channel walls, the corresponding displacement of the valve-opening center from the geometric channel centerline was obtained from

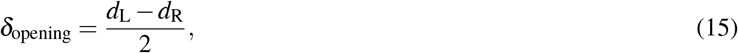

with

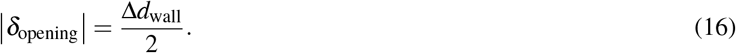

The bilateral clearance difference was used as the primary measure of geometric bias, while the opening-center displacement was retained as an equivalent geometric description. Image coordinates were converted to physical distances using the microscopy spatial calibration. Measurements were performed on the original image data; contrast adjustments used for figure preparation did not alter the coordinates used for geometric analysis.

### H. RBC-scale particle transport analysis

Finite-size particle transport was examined using spherical polystyrene particles with a nominal diameter of 8 *µ*m, providing an RBC-scale probe of post-valve transport and redistribution. The working suspension contained 1% 8 *µ*m polystyrene particles in deionised water supplemented with 23% density-gradient medium (CAS No. 92339-11-2; Sigma–Aldrich, Merck) to reduce gravitational settling. Particle experiments were performed in the rigid fixed-gap valve under the steady-flow conditions described in Section IV B. Image sequences were acquired using the optical system described in Section IV E at 30,000 frames s^*−*1^ with an exposure time of 4 *µ*s. Each recording contained approximately 2300–3000 consecutive frames, and the same downstream field of view was retained across all directly compared flow conditions.

Static channel and illumination features were removed by temporal-median background subtraction. Focused particles were identified using a custom image-analysis workflow based on the local optical response of the particle core and surrounding ring, with fixed detection settings used across the compared conditions. The centroid position of each accepted particle detection was recorded for every analyzed frame. A particle event was defined as one accepted centroid detection in one image frame.

Whole-field transport asymmetry was quantified from the cumulative centroid events recorded over each complete image sequence. Event counts were summed separately on the two sides of the geometric channel centerline and normalized by the corresponding valid analysis area before calculation of *AI*_event_ using Eq. (2). Each flow condition therefore yielded a single recording-level *AI*_event_ value based on the cumulative event-density distribution. Because the event map retains repeated detections of particles across successive frames, this metric characterizes the spatial distribution of particle-detection events within the analyzed field rather than the number of unique particles.

Particle passage through a fixed downstream section was evaluated separately to reduce the influence of repeated residence-time-weighted detections. Centroid detections in consecutive frames were linked into short particle trajectories using a nearest-neighbor criterion. Retained trajectories were required to contain at least three linked detections, and each trajectory was counted once when it crossed the prescribed downstream section at *x* = 450 *µ*m. Crossing events were assigned to the two sides of the channel centerline according to their interpolated lateral crossing position. The resulting crossing-count asymmetry was quantfied using *AI*_cross_ defined in Eq. (3). Thus, *AI*_cross_ reflects the bilateral distribution of particle passages through the downstream section, whereas *AI*_event_ describes the cumulative spatial distribution of detected particle events over the full analyzed field.

Downstream spatial redistribution was further assessed using two-dimensional centroid-event density maps constructed from all accepted detections within each recording. For direct comparison of the low- and high-Reynolds-number conditions, the event-density map obtained at the lowest *Re*_gap_ was subtracted from that obtained at the highest *Re*_gap_ after applying the same spatial registration, valid-area mask, and normalisation procedure. Positive and negative regions in the resulting high-minus-low-*Re*_gap_ map therefore represented relative enrichment and depletion of particle-detection events, respectively.

Particle-transport quantities were compared using the gap-based Reynolds number defined in Eq. (4), with the fixed rigid-valve opening. Detailed particle-detection criteria, trajectory-linking parameters, valid-area definition, event-density normalization, downstream-section coordinate conversion, and difference-map construction are provided in the Supplementary Methods.

Below is an alternative shorter version of section I

### I. Reproducibility and data summarisation

Because symmetry-breaking internal flows are sensitive to small geometric perturbations^11–13^, comparisons across flow conditions were performed within the same physical device for each valve configuration. The complete flow-rate series for the rigid fixed-gap, centered flexible, and geometrically biased flexible valves was therefore acquired using the same corresponding device, preserving the valve geometry, imaging field, and sampling coordinates while the imposed flow condition was varied. The whole-blood and RBC-scale particle measurements used the same valve configurations and provided cross-modality corroboration of the observed Reynolds-number-dependent behavior.

All devices were screened before quantitative acquisition to confirm intact bonding, unobstructed channels, functional valve structures, and the absence of visible leakage or other device dysfunction. For velocity-field measurements, PIVlab post-validation was used to remove clearly non-physical vectors using physically plausible velocity limits. The valid-vector percentage, defined as the proportion of reconstructed vectors retained after post-validation, ranged from 93–100% across the analyzed recordings. No routine manual exclusion of individual frames was applied.

For each recording, *AI*_mirror_ was calculated for every reconstructed velocity field. The highest and lowest 10% of frame-resolved values were removed, and the mean and standard deviation of the remaining central 80% were used as the condition-level value and its temporal variability, respectively. The resulting error bars therefore describe temporal frame-to-frame variability within a recording and not variability across independent devices. Particle-transport metrics were calculated at the recording level: *AI*_event_ from the bilateral distribution of all detected particle events over the analyzed sequence, and *AI*_cross_ from linked particle passages through the prescribed downstream section, as defined in Eqs. (2) and (3). These analyses were designed to resolve controlled within-device changes with flow condition and were not interpreted as estimates of population-level device-to-device variability.

## V. CONCLUSION

This study identifies a Reynolds-number-dependent symmetry-breaking transition as an organising feature of post-valve venous flow. The transition was resolved computationally and experimentally, was regulated by valve compliance and geometric bias, and remained detectable in whole blood. Once selected, the asymmetric flow organization also generated persistent heterogeneity in downstream RBC-scale transport.

Flow-state selection therefore provides an additional physical descriptor of valve-associated hemodynamics alongside conventional measures of velocity, shear, recirculation, and residence time. Its sensitivity to local inertia, valve mechanics, and small geometric perturbations provides a mechanism through which modest structural differences can generate substantially different downstream delivery and clearance environments. These findings establish post-valve symmetry breaking as a physically distinct contributor to spatially heterogeneous hemodynamics relevant to valve-associated thrombosis. The repeatable separation between the increasing- and decreasing-Reynolds-number branches under transient forcing further motivates controlled studies of its dynamical origin, transient relaxation, and dependence on prior flow evolution.

## SUPPLEMENTARY MATERIAL

See the supplementary material for additional computational, fluid–structure interaction, velocimetry, calibration, and particle-transport analysis details.

## ACKNOWLEDGMENTS

Y.C. acknowledges support from the University of Sydney Faculty of Engineering Research Scholarship.

## AUTHOR DECLARATIONS

### Ethics Approval

Non-identifiable human whole-blood samples were obtained from the Australian Red Cross Lifeblood Biological Resources program, which collected the samples with donor consent for research purposes. The University of Sydney Human Ethics Office determined that this lower-risk project was exempt from ethics review because it involved non-identifiable biospecimens collected with consent for research use.

### Author Contributions

Yuxin Chen: Conceptualization; Methodology; Investigation; Formal analysis; Visualization; Writing – original draft; Writing – review & editing.

Daniele Vigolo: Conceptualization; Methodology; Supervision; Resources; Funding acquisition; Writing – review & editing.

## DATA AVAILABILITY

The data that support the findings of this study are available from the corresponding author upon reasonable request.

