## Supplementary Information for "Symmetry-breaking flow transition as an under-recognized hemodynamic factor in valve-associated thrombosis"

### Supplementary Material for *Symmetry-breaking flow transition as an under-recognized hemodynamic factor in valve-associated thrombosis*

Yuxin Chen[1,2] and Daniele Vigolo[1,2]
[1)] *School of Biomedical Engineering, The University of Sydney, Sydney, New South Wales, Australia*
[2)] *Sydney Nano Institute, The University of Sydney, Sydney, New South Wales, Australia*
(*Electronic mail:)

#### I. CYCLE-RESOLVED ANALYSIS OF THE TRANSIENT SYMMETRY-BREAKING RESPONSE

To examine the repeatability of the branch separation observed in the geometrically symmetric rigid-valve model, the three consecutive forcing cycles were analyzed independently. Within the forward-flow phase of each cycle, the increasing- and decreasing-$Re_{\text{gap}}$ portions were identified from the temporal evolution of the instantaneous gap Reynolds number and retained as separate branches.

For each cycle, $AI_{\text{mirror}}$ was expressed as a function of $Re_{\text{gap}}$ separately for the increasing and decreasing branches. The three cycle-resolved branches were then linearly interpolated onto a common $Re_{\text{gap}}$ grid. Representative increasing and decreasing branches were obtained from the pointwise median across the three cycles,

$$\widetilde{AI}^{\uparrow}_{\text{mirror}}(Re) = \text{median}_{i=1,2,3}\left[AI^{\uparrow}_{\text{mirror},i}(Re)\right], \tag{1}$$

and

$$\widetilde{AI}^{\downarrow}_{\text{mirror}}(Re) = \text{median}_{i=1,2,3}\left[AI^{\downarrow}_{\text{mirror},i}(Re)\right], \tag{2}$$

where $i$ denotes the forcing cycle and the arrows denote increasing and decreasing $Re_{\text{gap}}$, respectively. The resulting median branches are those reported in Fig. 2B of the main manuscript.

The cycle-resolved trajectories showed consistent separation between the increasing- and decreasing-$Re_{\text{gap}}$ phases near the onset of pronounced asymmetry. The recurrence of this branch separation across successive forcing cycles supports the Fig. 2B. The full transient evolution of the geometrically symmetric rigid-valve model over all three forcing cycles is provided in Supplementary Movie S1.

The horizontal $AI_{\text{mirror}} = 0.05$ level shown in Fig. 2B was used as an operational reference for the onset region of pronounced asymmetry in the rigid-valve computational analysis. This reference level was specific to that analysis and was not used as a common threshold for the other computational or experimental datasets.

#### II. CONSTRUCTION OF THE RIGID- AND FLEXIBLE-VALVE RESPONSE CURVES

The rigid- and centered flexible-valve responses presented in Fig. 2C of the main manuscript were constructed from cycle-resolved $AI_{\text{mirror}}$–$Q$ trajectories. Within each forcing cycle, the increasing- and decreasing-$Q$ portions of the forward-flow phase were retained as separate branches.

For each valve configuration, the cycle-resolved branches were interpolated onto a common linearly spaced imposed-flow-rate grid. The corresponding $AI_{\text{mirror}}$ values were interpolated in logarithmic space. For each cycle,

$$A_i(Q) = \log_{10}\left[AI_{\text{mirror},i}(Q)\right], \tag{3}$$

where $i$ denotes the forcing cycle. Representative increasing and decreasing branches were obtained from the pointwise median across the three cycle-resolved curves,

$$\widetilde{A}^{\uparrow}(Q) = \text{median}_{i=1,2,3}\left[A^{\uparrow}_i(Q)\right], \tag{4}$$

and

$$\widetilde{A}^{\downarrow}(Q) = \text{median}_{i=1,2,3}\left[A_i^{\downarrow}(Q)\right]. \tag{5}$$

The representative branches were then transformed back to the original $AI_{\text{mirror}}$ scale according to

$$\widetilde{AI}_{\text{mirror}}(Q) = 10^{\widetilde{A}(Q)}. \tag{6}$$

The same procedure was applied independently to the rigid and centered flexible-valve datasets. In Fig. 2C, solid and dashed curves represent the median increasing- and decreasing-$Q$ branches, respectively, for each valve configuration.

#### III. COMPUTATIONAL FORCING AND FLEXIBLE-VALVE VISCOSITY PENALIZATION

The transient computational results presented in Fig. 2 of the main manuscript were generated using a periodic bidirectional forcing waveform. The elapsed time within each forcing cycle was defined as

$$t_c = \text{mod}\left(t, T_{\text{cycle}}\right), \qquad T_{\text{cycle}} = 3\ \text{s}. \tag{7}$$

Each cycle comprised a 2 s phase in one axial direction followed by a 1 s phase in the opposite direction. The target-velocity waveform was defined as

$$V_{\text{target}}(t_c) = \begin{cases} -V_{\text{peak}}\sin^2\left(\dfrac{\pi t_c}{2\ \text{s}}\right), & 0 \le t_c < 2\ \text{s}, \\ V_{\text{peak}}\sin^2\left[\dfrac{\pi\left(t_c - 2\ \text{s}\right)}{1\ \text{s}}\right], & 2\ \text{s} \le t_c < 3\ \text{s}, \end{cases} \tag{8}$$

with

$$V_{\text{peak}} = 0.117\ \text{m}\,\text{s}^{-1}. \tag{9}$$

The squared-sine waveform produced zero imposed velocity at the beginning of each cycle, at flow reversal, and at the end of the cycle, with smooth acceleration and deceleration within each directional phase.

The waveform was implemented through a pressure-driven inlet condition. For each computational geometry, the pressure amplitude was calibrated to reproduce a peak velocity of $0.117\ \text{m}\,\text{s}^{-1}$ in the corresponding reference flow condition. An initial transient simulation was performed using a trial pressure amplitude, the resulting peak velocity was measured, and the pressure amplitude was then rescaled proportionally before the final transient simulation. The same calibration procedure and temporal forcing profile were used for the rigid- and flexible-valve configurations.

For the flexible-valve simulations, a smooth viscosity-penalization treatment was applied as the leaflet opening approached closure. The local effective viscosity increased continuously according to the instantaneous valve opening and forcing phase, suppressing residual flow in the near-closure state. The baseline physical viscosity was retained in the calculation of the reported $Re_{\text{gap}}$.

#### IV. SUPPORTING DETAILS OF THE FLEXIBLE-VALVE FLUID–STRUCTURE INTERACTION MODEL

The compliance-dependent response presented in Fig. 2C of the main manuscript was obtained using a coupled fluid–structure interaction (FSI) model combining Laminar Flow, Solid Mechanics, and Moving Mesh in COMSOL Multiphysics 6.2. The fluid and leaflet domains were coupled along the fluid–solid interfaces, allowing the valve geometry to evolve continuously under transient hydrodynamic loading. The influence of flexible-valve mechanics on local venous-valve hemodynamics has been demonstrated previously[1].

The flexible leaflets were represented using the nearly incompressible two-parameter Mooney–Rivlin hyperelastic model specified in the main manuscript. The strain-energy density was expressed as

$$W = C_{10}\left(I_1 - 3\right) + C_{01}\left(I_2 - 3\right) + W_{\text{vol}}, \tag{10}$$

where $I_1$ and $I_2$ are the first and second invariants of the isochoric deformation tensor, and $W_{\mathrm{vol}}$ is the volumetric contribution associated with near-incompressibility. The constitutive parameters reported in the main manuscript were applied directly in the simulations.

For this formulation, the initial shear modulus is

$$G_0 = 2\left(C_{10} + C_{01}\right), \tag{11}$$

and, in the nearly incompressible small-strain limit,

$$E_0 \approx 3G_0. \tag{12}$$

The FSI simulations were governed directly by the Mooney–Rivlin constitutive parameters, while the equivalent Young's modulus provides a small-strain stiffness scale for interpretation of the leaflet material.

#### V. VELOCITY-FIELD ALIGNMENT AND MIRROR-ASYMMETRY ANALYSIS

Experimental velocity fields presented in Fig. 3A–B of the main manuscript were obtained using Ghost Particle Velocimetry (GPV), following established optical and image-analysis approaches for microfluidic velocimetry[2,3].

Non-fluid regions, including the valve structure and channel walls, were excluded using geometry-specific masks defined in PIVlab[4]. For the flexible-valve measurements, each flow condition was maintained for 5 min before image acquisition to allow the valve position and flow field to stabilize. Because the stabilized leaflet opening varied between conditions, the mask surrounding the valve region was adjusted to the corresponding geometry before velocity-field analysis.

The geometric reference for $AI_{\mathrm{mirror}}$ was defined from the bright-field image using a consistent set of predefined wall landmarks. Two landmarks were selected on each straight-channel wall and used to fit the upper and lower wall boundaries. The geometric channel centerline was defined as the midline between the two fitted wall boundaries. The same landmark locations were used across repeated analyses to maintain a consistent geometric reference.

The measured coordinates were transformed into a channel-aligned coordinate system, with $s$ directed along the fitted centerline and $n$ normal to it. The corresponding velocity components were denoted $U_s$ and $U_n$. Reflection across the geometric centerline was then defined as

$$\mathbf{u}_{\mathrm{mirror}}(s,n) = \left[U_s(s,-n), -U_n(s,-n)\right], \tag{13}$$

such that the channel-normal velocity component changes sign under reflection.

The analysis region extended to 75% of the measured channel half-width on either side of the centerline, while 3% of the axial extent was excluded at each end. The original velocity components and their reflected counterparts were interpolated onto a common $300 \times 160$ grid in $(s,n)$ coordinates using linear scattered interpolation without extrapolation.

Masked or non-finite velocity vectors were excluded before interpolation. Where vector-type information was available, masked vectors were excluded while regular and filtered or replaced vectors were retained. Following interpolation, only spatial locations containing finite velocity information for both the original and mirrored fields were retained for calculation of $AI_{\mathrm{mirror}}$.

#### VI. WHOLE-BLOOD VELOCITY-PLANE CALIBRATION

The velocity-plane conversion used for the whole-blood analysis in Fig. 3C(ii–iii) of the main manuscript was obtained from an independent two-plane calibration performed with water in the same rigid-valve geometry. Velocity was measured at the near-bottom and middle imaging planes over six imposed flow rates spanning 150–600 $\mu\mathrm{L\,min}^{-1}$. The corresponding relationships were fitted independently using linear regression, giving $R^2 = 0.99988$ for the middle-plane calibration and $R^2 = 0.99743$ for the near-bottom calibration:

$$U_{\mathrm{middle}} = 0.000811333\,Q - 0.000877964, \tag{14}$$

and

$$U_{\mathrm{bottom}} = 0.000352012\,Q - 0.0124179, \tag{15}$$

where $Q$ is the imposed flow rate in $\mu\mathrm{L\,min}^{-1}$ and velocity is expressed in $\mathrm{m\,s}^{-1}$.

The two fitted relationships were combined to obtain a direct conversion from the measured near-bottom velocity to the corresponding middle-plane velocity. From Eq. (15),

$$Q = \frac{U_{\text{bottom}} + 0.0124179}{0.000352012}, \tag{16}$$

which gives

$$U_{\text{middle}} = 0.000811333 \left( \frac{U_{\text{bottom}} + 0.0124179}{0.000352012} \right) - 0.000877964. \tag{17}$$

Equation (17) was applied to the near-bottom whole-blood velocity measurements before calculation of the corresponding $Re_{\text{gap}}$ reported in Fig. 3C(iii) of the main manuscript.

#### VII. SUPPORTING ANALYSIS OF RBC-SCALE PARTICLE TRANSPORT

Additional implementation details for the RBC-scale particle analyses presented in Fig. 5C(ii–iii) of the main manuscript are provided here.

Focused particles were identified using a ring-response detector designed for the bright-core/dark-ring appearance of the 8 $\mu$m particles in the high-speed image sequences. Candidate centers were identified from the local response

$$R = \left(I_{\text{core}} - I_{\text{ring}}\right) + 0.35 \left(I_{\text{outer}} - I_{\text{ring}}\right), \tag{18}$$

where $I_{\text{core}}$, $I_{\text{ring}}$, and $I_{\text{outer}}$ denote the mean image responses within the core, surrounding ring, and local outer region, respectively. The core radius was 1.7 pixels, the ring extended from 3.0 to 5.8 pixels, and the outer reference region extended from 7.0 to 11.0 pixels. Candidate peaks were required to exceed the 97.5th percentile of the positive ring response within the analyzed field and were separated by at least 5 pixels. Additional morphology criteria included a minimum dark-ring sector fraction of 0.35, a maximum ring coefficient of variation of 3.0, a minimum core circularity of 0.10, and a core area of 2–60 pixels$^2$. Contrast-, ring-darkness-, and edge-sharpness-based acceptance levels were determined automatically from a representative subset of frames using fixed quantile criteria and then applied to the corresponding image stack.

For the whole-field event-density analysis, a condition-specific downstream analysis region was defined to exclude structures outside the particle-carrying fluid domain. The local channel centerline divided this region into two sides. The analyzed pixel area on each side was calculated column by column relative to the local centerline, allowing differences in available analyzed area to be retained explicitly. The corresponding event densities were

$$\rho_{\text{event},1} = \frac{N_{\text{event},1}}{A_1}, \qquad \rho_{\text{event},2} = \frac{N_{\text{event},2}}{A_2}, \tag{19}$$

where $N_{\text{event}}$ is the accumulated number of accepted centroid events and $A$ is the valid analyzed area on the corresponding side. These area-normalized event densities were used to calculate $AI_{\text{event}}$ as defined in the main manuscript.

For comparison of the low- and high-flow spatial distributions, the centroid event-count maps obtained at 150 and 600 $\mu\mathrm{L\,min}^{-1}$ were normalized independently by their total event counts within the analyzed region,

$$M_Q^* = \frac{M_Q}{\sum_\Omega M_Q}, \tag{20}$$

where $M_Q$ is the event-count map at flow rate $Q$ and $\Omega$ denotes the analyzed region. The redistribution map was then calculated as

$$\Delta M = M_{600}^* - M_{150}^*. \tag{21}$$

Positive and negative values therefore represent relative enrichment and depletion, respectively, at 600 $\mu\mathrm{L\,min}^{-1}$ compared with 150 $\mu\mathrm{L\,min}^{-1}$. Gaussian smoothing with a standard deviation of 2 pixels was applied only for visualization of the spatial map; the quantitative subtraction was performed using the unsmoothed normalized event-count maps.

For the section-based analysis, the accepted centroid coordinates were linked into short trajectories by nearest-neighbor association. The maximum inter-frame linking distance was 18 $\mu$m, linking was restricted to detections in consecutive frames, and trajectories containing fewer than three detections were excluded. Tracks were additionally required to exhibit at least 2 $\mu$m net downstream displacement and to propagate in the positive downstream direction.

The crossing section was defined at a local position 150 $\mu$m downstream from the left boundary of the analyzed particle field. Its coordinate in the original image was obtained from

$$x_{\text{cross,global}} = x_{\text{FOV,left}} + 150\ \mu\text{m}, \tag{22}$$

with the pixel-to-distance conversion applied to the field-of-view origin. For the analysis reported in the main manuscript, this corresponded to the global downstream position $x = 450\ \mu$m.

For each linked trajectory crossing the prescribed section, the lateral crossing position was obtained by linear interpolation between the two trajectory points bracketing the section. Each trajectory contributed only its first accepted crossing. The crossing positions were centered relative to the channel field-of-view center, and the resulting upper- and lower-side crossing counts were used to calculate $AI_{\text{cross}}$ as defined in the main manuscript.

#### SUPPLEMENTARY MOVIE

**Supplementary Movie S1.** Time-resolved two-dimensional velocity field of the geometrically symmetric rigid-valve model over the full 9 s simulation, comprising three consecutive bidirectional forcing cycles. The post-valve jet selected the same lateral direction during the first two cycles and the opposite direction during the third cycle.
